# Simultaneous hybrid genome sequencing of *Vermamoeba vermiformis* and its Dependentiae endosymbiont *Vermiphilus pyriformis*

**DOI:** 10.1101/2021.04.23.440484

**Authors:** Vincent Delafont, Mégane Gasqué, Yann Héchard

## Abstract

A hybrid sequencing approach, using short and long reads sequencing, was employed for characterizing the genomes of the free-living amoeba host *Vermamoeba vermiformis*, along with its Dependentiae endosymbiont *Vermiphilus pyriformis*. The amoeba host reconstructed nuclear genome is 39.5 Mb, and its full mitochondrial genome is 61.7 kb. The closed, circular genome of the Dependentiae endosymbiont *Vermiphilus pyriformis*, naturally infecting *V. vermiformis* is 1.1 Mb.

## Rationale for the sequencing

Free-living amoebae are unicellular phagotrophic protists that are omnipresents in a wide range of environments (1). Among those, representatives of the species *Vermamoeba vermiformis* are recurrently found worldwide in natural and man-made ecosystems (2). Moreover *V. vermiformis* phagotrophic lifestyle encourages the establishment of transient and/or stable associations with numerous other microorganisms, including potential human pathogens, such as Legionella pneumophila and nontuberculous mycobacteria (3). Among those documented associations, recent work has put in evidence a stable, endosymbiotic association between an isolate of *V. vermiformis* and an endosymbiotic bacteria of the recently described Dependentiae phylum (4, 5). Such discovery enabled to access and maintain in laboratory culture one of the few Dependentiae isolates known to date. Having this culture at our disposal represented an opportunity to gather new knowledge on both microbial partners, for which genomic data remain very scarce to this day.

## Provenance, isolation, and growth conditions

The isolate *Vermamoeba vermiformis* TW EDP 1, naturally infected with the endosymbiont *Vermiphilus pyriformis*, was recovered from a tap water from Ivry/Seine, France, and has previously been described in details (4). *V. vermiformis* was recovered from a tap water sample filtered through a 3 µm nitrocellulosic membrane, which was deposited on a non-nutrient agar plate seeded with *E. coli* (6). The development of amoebae was regularly checked under a phase contrast microscope, and trophozoites cells were transferred in Page’s Amoeba Saline buffer (PAS; 4 mM MgSO_4_, 2.5 mM Na_2_HPO_4_, 2.5 mM KH_2_PO_4_, 0.4 mM CaCl_2_, 4 mM sodium citrate, pH 6.5) liquid cultures, supplemented with *E. coli* at a final concentration of approx. 5.10^8^ cells / mL. Liquid cultures were let to grow at 20°C until *E. coli* were cleared from the medium. Trophozoites were rinsed twice with PAS buffer to get rid of eventual remaining *E. coli* cells, and then detached using a cell scrapper. In total, two batches of approx. 2.10^7^ trophozoites were collected from three 175 cm^2^ culture flasks each, pelleted by centrifugation at 2’500 g for 20 minutes and stored at −20°C until DNA extraction.

## DNA extraction, library preparation and sequencing

One batch of pelleted cells was extracted using the Blood and Tissue kit (Qiagen), following manufacturer’s recommendations, using the QIAcube Connect apparatus (input material set as ‘bacterial pellet’, DNA was elution volume set at 150 µL). The resulting DNA extract was then sent for library preparation (Nextera XT, 8 PCR cycles) and short read sequencing (Illumina Nextseq 2000 platform) at the Microbial Genome Sequencing Center (https://www.migscenter.com/). The second batch was extracted using Gentra® Puregene® Yeast/Bacteria kit (Qiagen), following manufacturer’s recommendations for extracting DNA from cultured cells. DNA was eluted in a final volume of 50 µL. For Oxford Nanopore Technologies (ONT) sequencing, a library was prepared from 1.5 µg of purified DNA using the ligation sequencing kit (SQK-LSK109) following the ‘Genomic DNA by Ligation’ protocol (GDE_9063_v109_revX_14Aug2019 version). Library was loaded on an R10.3 flowcell, run for 24h on a MinIon Mk1C.

## Description of reads quality control and genome assembly

ONT sequencing generated 255’294 reads, representing 1’608’622’111 base pairs (N50 read length: 15,339 kb, median quality: Q11.3). Basecalling was performed using Guppy v4.0.11, using the ‘high accuracy” setting on the MinIon Mk1C. Illumina sequencing generated a total of 20’561’030 reads, representing 2’734’853’230 base pairs. Overall quality of reads was visualized using FastQC, and Illumina reads were subsequently trimmed at Q<30, using Cutadapt v 1.16 (7). Trimmed reads represented a total of 2’685’325’118 base pairs.

An initial assembly was performed using Flye v2.6 on ONT sequences only, with --nano-raw, --plasmids and --meta option enabled (8). Illumina reads were then mapped onto the ONT assembly using minimap2 (9). The produced bam file was further used for polishing and correcting the ONT assembly, using pilon v1.20, iteratively until no additional correction was done (10). The polished contigs were binned using Metabat2 v2.15, with a -minContig option set to 1500 (11). This step produced 4 bins, which were individually scrutinized by performing a taxonomic annotation of all contigs, using megablast algorithm (12, 13). Out of those bins, 2 comprised a collection of contigs with best BLAST hits matching with either Vermamoeba vermifomis, or other related eukaryotic microorganisms, and were thus merged to compose the genome assembly of *Vermamoeba vermiformis*. Another bin, comprising a single, circular contig was attributed to *Vermiphilus pyriformis*, while one bin comprised sequences best matching Escherichia coli (used as food source for the amoeba), and was discarded for further analyses.

## Genome characteristics of *Vermamoeba vermiformis* and *Vermiphilus pyriformis*

The sequencing of *Vermamoeba vermiformis* genome yielded 149 contigs, representing a total size of 39.5 Mb with an estimated completeness of 88.7% (Table 1). The genome size considerably differs from a previously published genome of another *V. vermiformis* strain (59.5 Mb), though GC content is comparable (14). Among contigs attributed to V. vermiformis, one stood out by its high coverage and low GC content, as compared with other *V. vermiformis* contigs. This highly covered contig, representing 67.1 kb, corresponded to the mitochondrial genome of *V. vermiformis* (Table1).

**Table 1:** statistics and features of *V. vermiformis* and V. pyriformis genome assemblies.

|  | *Vermamoeba vermiformis* | *V. vermiformis* mitochondrion | *Vermiphilus pyriformis* |
| --- | --- | --- | --- |
| N° of contigs | 149 | 1 | 1 |
| Total length | 39'458'332 | 67'148 | 1'123'447 |
| Coverage | 32x (Illumina) / 23x (ONT) | 1069x (Illumina) / 657x (ONT) | 959x (Illumina) / 346x (ONT) |
| GC content | 42.53% | 31.58% | 36.44% |
| Largest | 945'495 | 67'148 | 1'123'447 |
| N50 | 527'084 | 67'148 | 1'123'447 |
| N75 | 436'737 | 67'148 | 1'123'447 |
| L50 | 32 | 1 | 1 |
| L75 | 53 | 1 | 1 |
| Genome completeness | 88.7%<sup>1</sup> | NA | 92.8%<sup>2</sup> |
| Genome contamination | 2.4%<sup>1</sup> | NA | 0% |

**1** Genome completeness was estimated for *V. vermiformis*, using BUSCO v4.1.2 and the Eukaryota the lineage. Contamination represents the number of duplicated gene and is therefore speculative (16).
**2** Genome completeness was estimated for *V. pyriformis*, using CheckM v1.0.18 (17). The systematic absence of 4 marker from CheckM single copy marker genes has been taken into account for completeness calculation (5).

The sequencing effort conducted in this study also allowed for the complete reconstitution the Dependentiae endosymbiont *Vermiphilus pyriformis* circular genome. Thus, V. pyriformis was estimated to represent 1.1 Mb, with a GC content of 36.44%. Those features are highly similar to the closest relative of V. pyriformis with a sequenced genome (4, 15).

## Data availability

ONT and Illumina raw reads were deposited on the Sequence Read Archive under accession numbers <u>SRR14181580</u> and <u>SRR14181581</u>, respectively. All presented genome assemblies are available using Bioproject number <u>PRJNA720556</u>, at the NCBI.

## Acknowledgments

This work was partly supported by the region Nouvelle Aquitaine and Europe through the Habisan CPER-FEDER program, and by investment grants from the University of Poitiers.

